# Primary pterygium was not associated with corneal endothelial cell decrease in a rural Chinese population

**DOI:** 10.1101/595892

**Authors:** Yijun Hu, Li Li, Wei Qi, Zhenhao Liu, Yingpeng Liu, Ling Yuan

## Abstract

**Purpose:** To investigate the corneal endothelial cell density (ECD) in eyes with primary pterygium.

**Methods:** We first conducted a retrospective study to compare the ECD between 1670 eyes with primary pterygium and 4060 eyes without pterygium. Then we designed a prospective study to confirm the critical findings of the retrospective study in 95 patients with unilateral primary pterygium.

**Results:** In the retrospective study, the mean preoperative ECD in eyes with primary ptergyium (2453±306 cells/mm^2^) was statistically significantly lower than those in eyes without pterygium (2529±313 cells/mm^2^, *P*<0.0001). However, the difference was minor (76 cells/mm^2^, 3.1%). In the prospective study, the mean ECD in eyes with unilateral primary pterygium (2480±263 cells/mm^2^) was not significantly different from those in the contralateral eyes (2527±277 cells/mm^2^, *P*=0.20). The hexagonality (*P*=0.10) or coefficient of variation of size (*P*=0.15) of corneal endothelial cells was not significantly different between eyes with pterygium and the contralateral eyes.

**Conclusion:** Primary pterygium may not be associated with ECD decrease in our study cohort.

## Introduction

Pterygium is an ocular surface disorder which involves growth of the bulbar conjunctiva onto the cornea [1]. Over-exposure to ultraviolet light is the most significant risk factor associated with pterygium development and people living at the tropical areas or working outdoor are more likely to have the disease [2–4]. Studies have shown that apart from causing corneal surface changes [5–6], pterygium may also be associated with degradation of the Bowman’s layer and stromal scarring [7–8] which may attribute to local inflammatory and angiogenesis mediators [9]. Theoretically, these mediators and ultraviolet light may also damage corneal endothelium cells (ECC) and corneal endothelial cell density (ECD) may be decreased in eyes with pterygium [10]. Although results from previous clinical studies were consistent with this theory [11–13], in our cataract clinic we found that the preoperative ECD was not significantly decreased in cataract eyes with primary pterygium compared to those without pterygium. We speculated that pterygium might not be associated with ECD decrease in our population. To confirm our speculation, we reviewed the medical records of eyes underwent cataract surgery in our hospital and compared the preoperative ECD between 1670 eyes with primary pterygium and 4060 eyes without pterygium. We found that ECD decrease in eyes with pterygium was minor and it might attribute to the variation of ECD measurement. To confirm our speculation we designed a prospective study to determine the variation of ECD measurement and compare the ECD between the pterygium eyes and the contralateral eyes in 95 patients with unilateral primary pterygium. We found that the ECD was not significantly decreased in eyes with pterygium in our study cohort.

## Materials and Methods

The studies have been approved by the Institutional Review Board and was in agreement with the Declaration of Helsinki. In the retrospective study we reviewed the medical records and database recording ECD of eyes underwent cataract (from January, 2014 to September, 2016), and consecutively included all eligible eyes (1670 eyes with pterygium and 4060 eyes without pterygium) for analysis. Inclusion criteria was untreated primary pterygium and the patients age ≥51 years old. To prevent possible co-existing corneal endothelium pathology and ECD measuring bias, we only included eyes with a preoperative ECD of 1800-3500 cells/mm^2^ [14]. Eyes with pseudopterygium, recurrent pterygium, corneal dystrophy or corneal degeneration, history of corneal infection, glaucoma, uveitis, ocular trauma or intraocular surgery were excluded. We calculated the sample size of the prospective study according to the ECD in the retrospective study. A sample size of 95 patients would have 80% power to detect 5% difference in ECD at the confidence level of 95%. Therefore, in the prospective study we recruited 95 patients with untreated unilateral primary pterygium (from August, 2016 to June, 2017). Inclusion criteria exclusion criteria were the same as the retrospective study except there was no age limitation. Informed consent was obtained from each patient. The authors had access to information that could identify individual participants during or after data collection, but no individual participants could be identified according to the data presented in this article. All of the eyes underwent routine preoperative examination including best-corrected visual acuity (BCVA), intraocular pressure (IOP), anterior segment examination by slit-lamp, A-scan and B-scan ultrasonography (in the retrospective study). ECD was measured by the same technician using a specular microscope (SP-3000P; Topcon) and the center-to-center method [11]. ECD was measured one time for each eye with a cell count of at least 60 cells in the retrospective study and was measured three times consecutively for each eye with the cell counts of at least 80 cells in the prospective study. In the prospective study, the coefficient of variation (COV) and SD of the three consecutive ECD measurements were calculated. The average of three consecutive ECC parameters including ECD, hexagonality and coefficient of variation of size (ECV) from each eye was used for analysis. Data was presented as mean ± standard deviation (SD). Two-tailed Student’s *t*-test was used for the comparison of two sets of normally distributed data and Mann-Whitney test was used otherwise. One-way ANOVA was used to compare three or more sets of normally distributed data and Kruskal-Wallis test was used otherwise. *P*<0.05 was considered statistically significant.

## Results

### Basic characteristics

In the retrospective study there were 1670 eyes with pterygium (PT group) and 4060 eyes without pterygium (NPT group). Among the patients who had received cataract surgery in both eyes within one year, 98 patients had bilateral pterygium, 150 patients had unilateral pterygium and 398 patients had no pterygium. There was significant difference in age between patients in the PT group (71.4±8.1 years) and those in the NPT group (70.5±8.3 years, *P*=0.0002), but not among patients with bilateral pterygium (70.3±6.9 years), unilateral pterygium (70.1±8.1 years) or without pterygium (68.6±8.2 years, *P*=0.10). Gender distribution was not significantly different between patients in the PT group and those in the NPT group, or among patients with bilateral pterygium, unilateral pterygium or without pterygium (*P*>0.05).

The mean age of the patients (40 males and 55 females) in the prospective study was 63.7±8.8 years. Forty-nine patients had pterygium in the right eyes and 45 patients had pterygium in the left eye.

### ECD in the retrospective study

The mean preoperative ECD in the PT group (2453±306 cells/mm^2^) was statistically significantly lower than those in the NPT group (2529±313 cells/mm^2^, *P*<0.0001). The difference in ECD between the PT group and the NPT group was 76 cells/mm^2^ (3.1%). There was statistically significant difference in preoperative ECD among patients with bilateral pterygium (2444±329 cells/mm^2^), unilateral pterygium (2486±309 cells/mm^2^) or without pterygium (2535±303 cells/mm^2^, *P*=0.001). In subgroup analysis, patients with bilateral pterygium had lower ECD than patients without pterygium (91 cells/mm^2^, 3.7%, *P*=0.01). However, in patients with unilateral pterygium, the ECD of the pterygium eyes (2487±313 cells/mm^2^) was very similar to the ECD of the contralateral eyes (2486±306 cells/mm^2^, *P*=0.99). The similar ECD in both eyes of patients with unilateral pterygium raised our suspect that ECD might not be decreased in eyes with pterygium in our population. We speculated that the small difference in ECD between the PT group and the NPT group might have been due to the variation of ECD measurement. The speculation was tested and confirmed in the prospective study.

### ECD in the prospective study

In the prospective study, the intraclass correlation coefficient (ICC) of single ECD measurement was 0.876, suggesting good reliability of the measurement. The 95% confidence interval (CI) was 3.0%-3.7% for the ECD COV, and was 76-91 cells/mm^2^ for the ECD SD.

In the prospective study, the average of three ECD measurements was 2480±263 cells/mm^2^ in the pterygium eyes and was 2527±277 cells/mm^2^ in the contralateral eyes (*P*=0.20). The difference in ECD between the PT group and the NPT group (76 cells/mm^2^, 3.1%) was just within the 95% CI of the ECD SD and COV, which had proven our speculation.

The ECC hexagonality was not significantly different between the pterygium eyes (52.8%±7.1%) and the contralateral eyes (51.4%±6.9%, *P*=0.10). The ECV was also similar between the pterygium eyes (36.3%±4.2%) and the contralateral eyes (37.0%±4.6%, *P*=0.15).

## Discussion

Pterygium is commonly seen in areas within the “pterygium zone” and in people with outdoor occupations [1–4]. In our retrospective study, 29.1% (1670/5730) of eyes underwent cataract surgery had concomitant pterygium. Pterygium may cause corneal surface irregularity and astigmatism [5–6]. Previous studies had shown that pterygium might also be associated with deeper corneal changes [11–13]. Mootha et al. first described changes at the corneal endothelium and the Descemet membrane underlying or directly adjacent to the pterygium [13]. In their study the mean ECD decrease was 367 cells/mm^2^ in eyes with unilateral pterygium [13]. Hsu et al. also detected an ECD decrease of 230 cells/mm^2^ (9.75%) in eyes with unilateral pterygium compared to the contralateral eyes [11]. They showed that 48.9% of the patients with unilateral pterygium had significant ECD decrease in the affected eyes, and the extent of ECD decrease was correlated with the size of the pterygium [11]. However, the results of our study came to a different conclusion. The ECD decrease in eyes with pterygium from our retrospective study was far less (76 cells/mm^2^, 3.1%) than those in the previous studies [11, 13, 15]. In our retrospective study, the ECD of eyes with unilateral pterygium was almost the same as the ECD of the contralateral eyes. In the prospective study, ECC hexagonality, ECV in eyes with primary unilateral pterygium was not significantly different from those in the contralateral eyes. The mean difference in ECD between pterygium eyes and the contralateral eyes was only 46 cells/mm^2^ (1.9%). Moreover, we confirmed that the ECD decrease in the retrospective study was due to the variation of ECD measurement.

The repeatability of ECD measurement by specular microscope is largely dependent on the cell count measured [16]. Doughty et al. found that a cell count of 25 cells could result in ±10% of ECD variation while an ECD variation of ±2% could be achieved if 75 cells were measured [16]. In our prospective study, a cell count of at least 80 cells was used and the 95% CI was 3.0%-3.7% for the ECD COV, and was 76-91 cells/mm^2^ for the ECD SD. These results were similar with those in the study of Doughty et al [16]. In our retrospective study the cell counts were at least 60 cells and the variation of ECD measurement would be expected to be larger. Moreover, the age difference between the PT group and the NPT group in the retrospective study was 0.9 year. The age-related ECD decrease in the Chinese of our patients’ age group was 0.16%/year [17]. Taking together we believed that primary pterygium might not be associated with ECD decrease in our study population.

The mechanisms of ECD decrease in eyes with pterygium are not fully understood. Two possible mechanisms have been speculated. One of them is the damage caused by ultraviolet light. This is based on the fact that ultraviolet light over-exposure is considered to be the most significant risk factor of pterygium [2–4], and ultraviolet radiation may trigger inflammation, oxidative damage and apoptosis of the ECC and cause ECC loss [10, 18]. In a previous study, outdoor workers with more UV radiation exposure was shown to have lower ECD compared to non-outdoor workers [19]. In a recent study, ECD decrease in eyes with pterygium was also found to be associated with the duration of UV radiation exposure [15]. The other cause of ECD decrease in eyes with pterygium is a variety of pathogenic factors including inflammation, angiogenesis and extracellular matrix modulators [9]. These modulators have been shown to be expressed in pterygium tissue or the cornea underlying or adjacent to the pterygium [7, 20]. Moreover, the levels of these pathogenic factors may be correlated with the clinical features of the pterygium [21–22].

There might be several reasons why primary pterygium was not shown to be associated with ECD decrease in our study. Firstly, the UV exposure in our patients might not reach to a certain threshold to cause ECD decrease. Li X et al. divided the patients with unilateral primary pterygium into two groups according to their daily UV exposure. In patients with longer UV exposure, the ECD in the eyes with pterygium was significantly lower than those in the contralateral eyes. However, in patients with short UV exposure the ECD was not significantly different between the eyes with pterygium and the contralateral eyes. Moreover, patients with longer UV exposure had significantly lower ECD than patients with short UV exposure [15]. It appeared that in eyes with pterygium the UV exposure needed to reach a certain threshold to induce ECD decrease. In our study, we did not record the UV exposure of the patients. Therefore, it was possible that the mixture of patients with short UV exposure had caused the insignificant decrease of ECD in eyes with pterygium. Secondly, the eyes in our study might had different clinical characteristics of the pterygium from those in the previous studies. Mootha et al. had shown that ECD decrease was found in pterygium with long duration [13]. Moreover, pterygium fleshiness, redness and extent may be correlated with the levels of inflammation, angiogenesis and extracellular matrix modulators which are involved in ECD decrease in eyes with pterygium [9]. Pterygium with intermediate fleshiness was shown had higher expression of extracellular matrix modulators [21]. Pterygium with intermediate redness and extent were also shown to have higher levels of chronic inflammation in the corneal stroma [22]. The eyes in our studies might have relatively small and less fresh pterygium since cataract surgery was indicated to these eyes. Thirdly, UV light radiation might have caused similar extent of ECD decrease in both eyes of our patients. This could be supported by the fact that in both our retrospective study and prospective study, ECD in eyes with unilateral pterygium was not significantly different from those in the contralateral eyes. Fourthly, pterygium might not cause significant ECD decrease in our study cohort, which might be due to adaption of the cornea endothelial cells to UV light radiation and chronic stromal inflammation.

In conclusion, the results of our studies suggested that primary pterygium might not be associated with ECD decrease in our study cohort.

## Acknowledgements

The study was supported by the Science Research Foundation of Aier Eye Hospital Group (AF2018003) and the Science and Technology Planning Project of Shanwei City (2017C015).

